# Robust Cluster Typing, a computationally accessible tool for species designation from whole genome data, exemplified with the genus *Campylobacter*

**DOI:** 10.64898/2026.11.17.765926

**Authors:** Lisa C Crossman, Claire Hill, Arif M Tanmoy, Nikkon Sarker, Hafizur Rahman, Heather Felgate, Nicol Janecko, Bilal Djeghout, Amira Roess, Senjuti Saha, John Wain

**Affiliations:** University of East Anglia, Norwich Research Park, Norwich NR4 7TJ, United Kingdom; SequenceAnalysis.co.uk, Norwich Research Park, Norwich NR4 7UG, United Kingdom; Centre for Microbial Interactions, Norwich Research Park, Norwich NR4 7UG, United Kingdom; Microbes and Food Safety, Quadram Institute Bioscience, Norwich Research Park, Norwich NR4 7UQ, United Kingdom; Child Health Research Foundation, SEL Huq Skypark, 23/2 Khilji Road, Block-B, Mohammadpur, Dhaka-1207, Bangladesh; George Mason University, College of Public Health, 4400 University Drive, Fairfax, Virginia 22030

**Keywords:** Robust clustering, bacterial typing, species assignment, *Campylobacter*, tree of life

## Abstract

Bacterial genomic analysis is a cornerstone of modern epidemiology. Many genomic typing methods exist, and most are designed to work within a given species in a hierarchical fashion. However, designation of bacterial species can be problematic, making the higher resolution level of hierarchical typing difficult. Using the universally present ribosomal proteins, we have developed a pipeline for a robust typing system which can be used to assign an unknown organism to a species-level cluster or, sometimes, a sub-species cluster. As proof of principle, we have used bacterial genomes from the genus *Campylobacter*, the most common cause of human bacterial gastroenteritis within which new strains, and new species (*C. infantis*), are reported. Species assignment, of all but the two most common species *C. jejuni* and *C. coli*, is beyond the ability of most microbiology laboratories.

Outbreak investigation, even with genomic data, is difficult and so a robust typing scheme would be useful.

Using available sequence-based typing schemes, we attempted to differentiate between 39 published *Campylobacter* species; we determined that the ribosomal 16S sequence typing and SNP typing did not discriminate all 39 but a single gene phylogeny method (*gyrB*), BUSCO, Mash, Panaroo and use of the Hug proteins differentiated all 39. We assessed computational accessibility (important for implementation) and biological applicability (one universal typing scheme is easier to validate). The application of Hug proteins proved computationally efficient and should be widely applicable. We developed a workflow in snakemake and Python, capable of clustering thousands of bacterial genomes. This Robust Cluster (RC) typing workflow was made accessible via the Terra platform, with usage instructions provided through an online training course on YouTube. Ninety-eight *Campylobacter* isolates from Bangladesh were sequenced and analysed using the Terra workflow, resulting in the identification of several novel clusters not previously found among publicly available genomes. Our work establishes the availability of RC typing and demonstrates its capacity to identify new *Campylobacter* lineages, thereby facilitating reporting and supporting future research initiatives.

## Introduction

Species designation is fundamental to our understanding of biological diversity. Bacteria within a species share genetic content, physiological traits and ecological niche. They are usually subdivided into smaller clusters, strains, for a more detailed analysis. However, naming bacterial isolates at the species level, even with DNA sequence data, can be problematic because the level of phenotypic and genotypic variation used to designate a species classification of bacteria is not consistent; rather it depends upon expert opinion (1). Taxonomy, however, aims to classify organisms to allow identification that is stable, and genomic guidelines have been suggested to promote the robust naming of new bacterial groups (2). The release of databases, such as the Genome Taxonomy Database (3), has revolutionised our ability to employ approaches combining average nucleotide identity (ANI) and relative evolutionary divergence for the investigation of bacterial clades – but to investigate diversity we need to populate these databases with more data from less well sampled parts of the bacterial population. One barrier to this is that the explosion in computational tools, for the analysis of whole genome sequence (WGS) datasets (4), makes choice difficult for the non-specialist.

The choice of analysis depends on the task in hand and the computational power available. For example, identification of a bacterial species for reporting needs a comprehensive set of reference genomes for comparison (5), whereas outbreak investigation or ecological guild definition requires higher resolution phylogenomic clustering and the incorporation of metadata (6). Hierarchical clustering is a common and effective approach which helps manage increasingly complex datasets (7) but the increasing sophistication introduces the risk of limiting accessibility through restrictive computational demands. Generating sequence data is now easier than analysing it. A balance must be struck between accessibility, cost, rigour, reproducibility and the granularity of subdivision when analysing genomic data (5).

Most high-resolution typing schemes are developed for a single species and so species designation generally provides the baseline of most hierarchical clustering algorithms; a simple, robust method for clustering genomes at the species level would enable laboratories generating sequence data to rapidly exploit their own results through direct reporting, followed by deposition into global databases to facilitate higher resolution typing to drive biological investigations. Campylobacteriosis represents a challenge to outbreak investigations by public health laboratories (8) and for community-based research projects (9).

*Campylobacter* is the most common cause of bacterial gastroenteritis globally (more common in developing countries) and the most common cause of food poisoning in most developed countries, yet our ability to type novel strains or recognise new species is limited (10). Outbreak investigation using WGS can be useful for *C. jejuni* (8) but very little is known about the other species (11). Sequence typing and core genome comparisons are the most successful genomic typing methods currently used, including CampyGStyper (12), but these methods are computationally demanding, often species based, not available to all laboratories and are not validated for national surveillance. A universal scheme for *Campylobacter* typing would go some way to resolving these issues, especially if applicable to all bacteria. Several universal methods exist for assigning species to bacterial genomes, e.g.: rMLST (13) 16S phylogenies (Supplementary Figure S1) (14), (single gene (*e.g. gyrB*) phylogenies (15), BUSCO (16), ribosomal proteins (17) and Mash, kmer distances (18). These methods all cluster organisms with a similar DNA sequence of a shared genomic feature (core, or coding sequence) into groups for which a specific scheme can be developed for more granular typing (19). The use of WGS data for the typing of other bacterial pathogens is facilitated by the existence of genomic methods which can reproduce traditional methods, *e.g.* geno-serotyping (20) allows validation and adoption of WGS-based typing by public health authorities for Salmonella (21). However, for *Campylobacter* this is not so straightforward, traditional typing has not been as successful (22) and there is no fast track to a validated genomic typing scheme. Also individuals may harbour more than one strain (23) and the source (or transmission vehicle) may harbour more than one strain (9), so multiple isolates must be analysed. These difficulties have led to a decline in the submission of isolates to reference laboratories for typing in developed countries (24). In developing countries the situation is worse (25) and the majority of *Campylobacter* infections, which occur in resource-poor regions of the world, are rarely sequenced or submitted to databases (26); this limits the global benefit from the use of genomic databases and hinders studies of bacterial diversity in particular. Whilst database independent methods of multi-locus sequence typing (27–29) exist for *Campylobacter* they fail to include all species (30) and/or require a level of compute power which limits their use on large datasets. A universal, accessible typing scheme would go some way to addressing these issues.

Here we compare several methods for typing *Campylobacter*, we use discrimination power (compared to the BUSCO method) (31) and required compute power as criteria for selection. The first step in improving *Campylobacter* genomic surveillance is to enhance database coverage to include more bacterial genomes from less well-resourced regions of the world. We hope that this method will facilitate that process.

## Results

Our aim was to facilitate the submission of more diverse genomes to public databases by determining which clustering tools would be suitable for use by non-professional informaticians in resource-poor research facilities, specifically in low to middle income countries. We tested the tools in three stages of increasing complexity and then adapted, and made accessible, the best for use in a low-compute power environment.

### Comparison and testing of *Campylobacter* genus-wide clustering methods

Three separate datasets were used to test the level of discrimination seen with the most used typing tools in a staged benchmarking:

Pre-testing preparation – Generation of a ribosomal sequence dataset for the genus (*Campylobacter*) under investigation – *Campylobacter* Ribosomal Sequence set from a reference genome

Stage one – Species assignment: use of 39 representative species of *Campylobacter* (32). Typing methods were compared for discriminatory level and computational intensity; the database used recognised species’ genomes from ref 32 and the result generated benchmarking of methods (Table 1)

**Table 1:** Benchmarking of *Campylobacter* species assignment using available methods on 39 *Campylobacter* genomes representing different species, *gyrB* and RC protein were the fastest to define all 39 groups.

| Method | Time to completion | Number of non-identical groups |
|---|---|---|
| Single gene (*gyrB*) | 19.771 +/- 3.402 s | 39 |
| Hug/RC nt | 58.124 +/- 1.308 s | 39 |
| Hug/RC protein | 25.440 +/- 0.179 s | 39 |
| BUSCO | 8 min 10s<sup>\$</sup> | 39 |
| Mash | 15.87 +/- 0.175 s | 39* |
| CheckM | 36 min 34s | 39 (2 lineage divisions) |
| Panaroo<sup>\$</sup> | 5 hr 1 min 46s | 39 |
| PopPUNK lineage model | 2.538 +/- 0.062 s | 7 lineages |

\*Distance estimation only
<sup>\$</sup>Alignment process took much of the time due to sequence length

Stage two – Population clustering: use of all publicly available, quality screened, genomes of *Campylobacter* [accessed October 2022] to generate a cluster reference set using the best method from Stage 1. Terminology: *Campylobacter* global dataset (all publicly available genome sequences annotated as *Campylobacter*); Stage 2 result generated the Cluster Reference Set (683 genomes used to define clusters for this genus.)

Stage three – Investigating isolates from Bangladesh: 98 genomes of strains isolated in Bangladesh were clustered using the Cluster Reference Set.

### Stage one: Species assignment (Table)

We compared for accuracy and useability the common methods used for assigning the species of *Campylobacter* to a genome.

### Stage 2 – clustering of all publicly available genomes of *Campylobacter*

For the *gyrB* method, we used the nucleotide sequences. For ribosomal protein sequences (Ribosomal Sequences Set, RCprot), we also used protein sequences. Although sequence conservation across kingdoms is well described (33), we observed considerable variation in protein length (Figure 1). We therefore introduced a quality control procedure that only included proteins within 1.5x the interquartile range (IQR) of length for all accessible *Campylobacter* protein sequences. We selected RC sequences from *C. jejuni* NCTC 11168 as the comparator.

**Figure 1.**
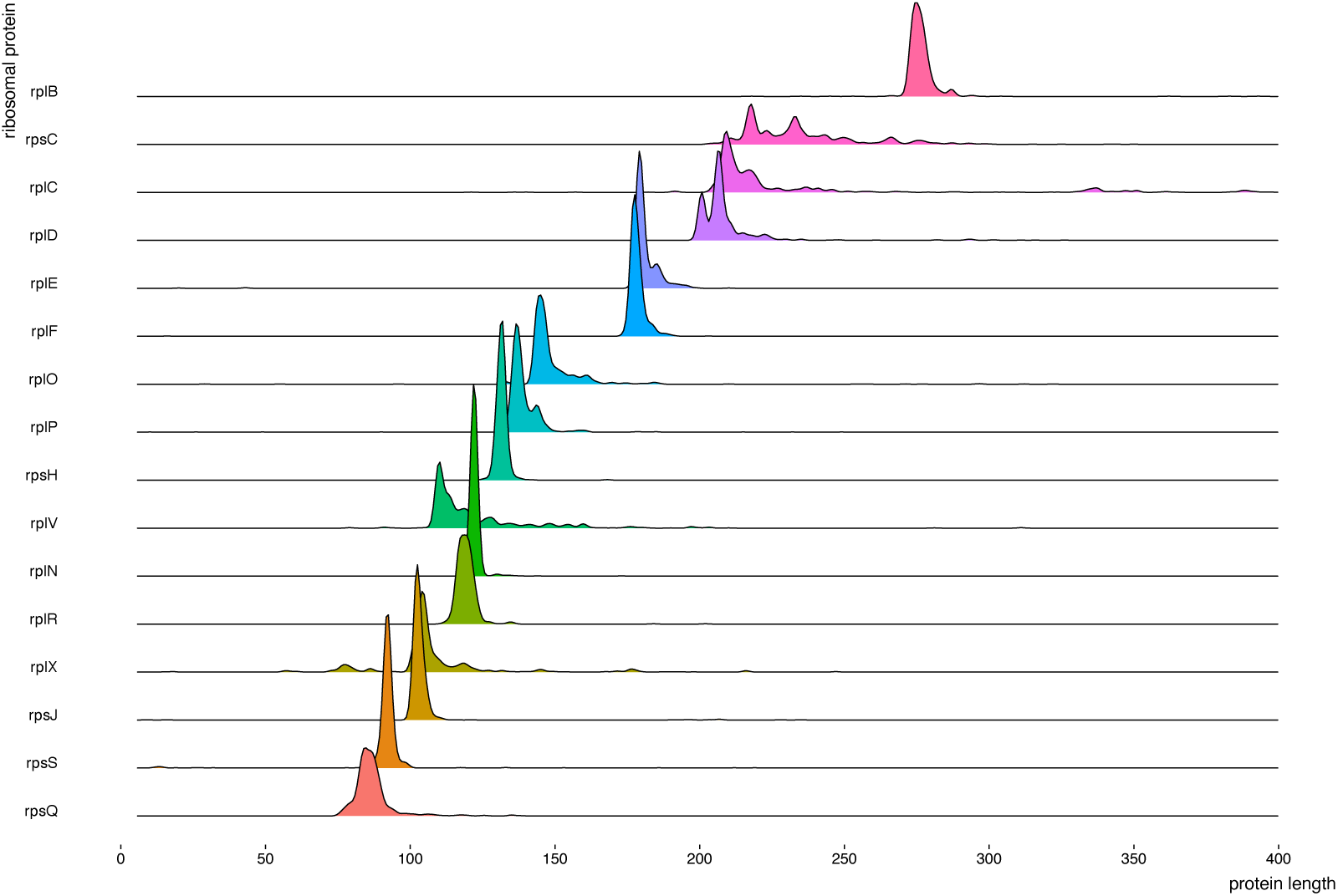
Conservation of the ribosomal proteins by length across curated Uniprot proteins. A dataset of all the manually curated Uniprot ribosomal proteins across all kingdoms was downloaded and the lengths of each determined and plotted. Some of the ribosomal proteins exhibit diverse protein lengths, while others have a more conserved length.

### Robust clustering workflow

The sequences of 16 ribosomal proteins conserved across all kingdoms of life (17, 34) were used in a workflow written in snakemake (35) and Python allowing the user to study these proteins in a set of genomes of interest by generating a phylogeny for use in high-level clustering.

The full workflow included the main workflow supported by two subworkflows (Figure 2), one with the ability to download reference genomes and gather GenBank files as well as conversion to nucleotide DNA sequence or protein fasta sequences. Both workflows were successfully accessed by our biologists from Terra.bio using JIDC Dockstore (see link in code availability section). The main workflow uses a downloaded Ribosomal Sequence Set collected from whole genome sequences.

**Figure 2.**
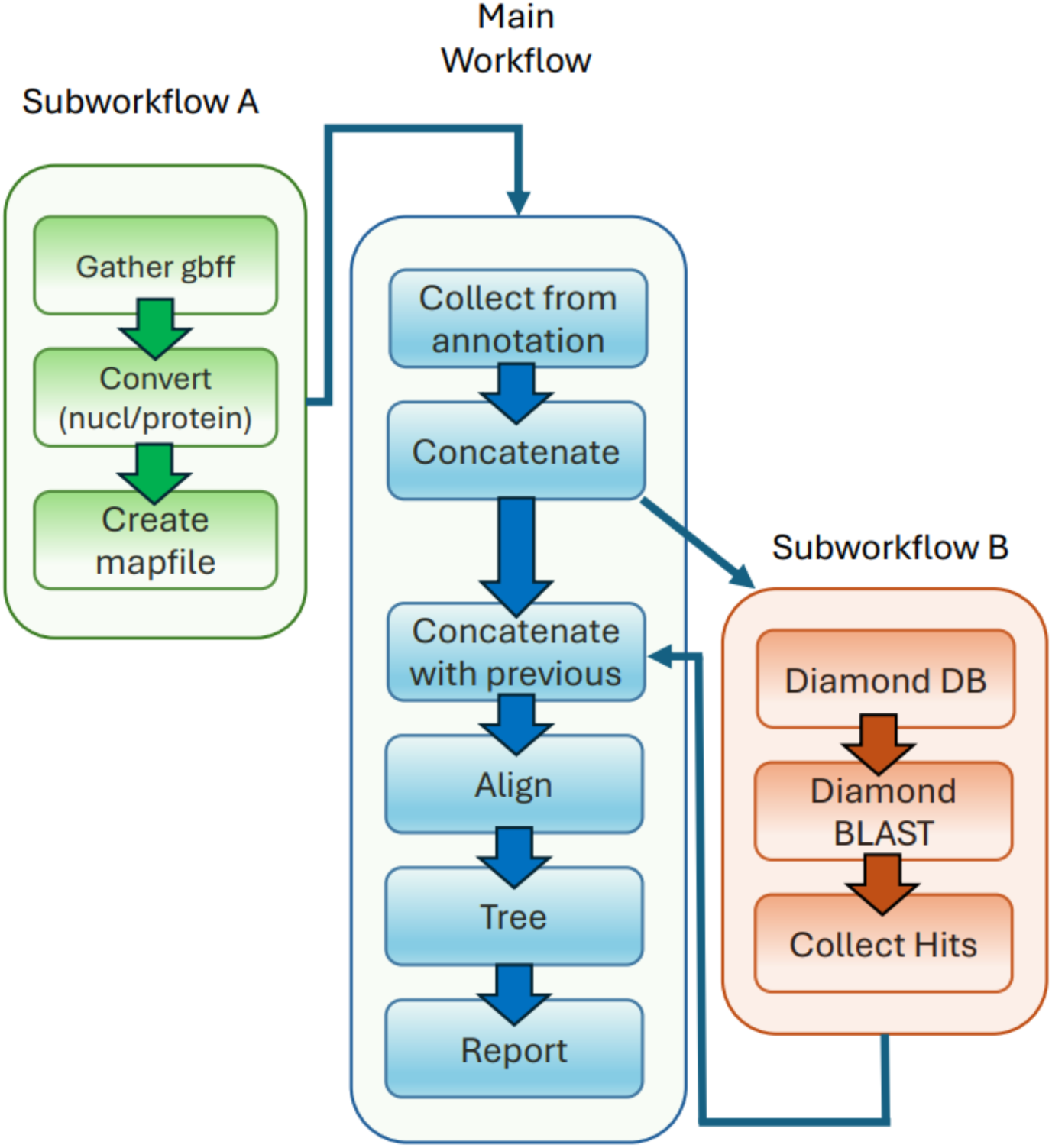
A snakemake workflow for RC clustering of whole genome sequences. Subworkflow A (green) is employed to gather whole genome sequence files as GenBank files, create nucleotide or protein fasta files dependent on the chosen run parameters and create a mapfile of GenBank filenames to organism names from the DEFINITION line of the GenBank file. In the main workflow (blue), sequences can be collected from annotation directly and concatenated into a form appropriate for fast alignments. If sequences are found to be missing in the annotation, subworkflow B (orange) is run to identify and extract those sequences using BLAST. At this stage, the main workflow concatenates all sequences, including any requested from previous runs. The concatenated sequences are aligned using Mafft, Trees are created using either Iqtree2 or Fasttree as requested in the chosen parameters and results are finalised. Training material is available at https://www.youtube.com/playlist?list=PLU47xRg_MKJpg8roYd9jTI-qOTHukGxcC

Gene (RCdna) or predicted protein (RCprot) sequences can be selected depending on whether DNA or protein has been chosen. In the second subworkflow, if any of the sequences were missing, after recovery from the annotated GenBank format files, these were searched for using BLAST. Sequences were discarded unless all 16 sequences were present following this step. All sequences were then concatenated together, aligned and a phylogeny created.

### Clustering the alignment

We tested several clustering methods for processing the RC pipeline output, including hierarchical clustering and machine learning. Ultimately, we selected MMSeqs2 (36) because it will generate clusters containing a single genome, using sequence files at 100% sequence identity, and is scalable to very large datasets. MMSeqs2 was used to generate a set of RC sequences that contained a single representative sequence from each cluster when all publicly available sequences were analysed, this set is called the *Campylobacter* Cluster Reference Set.

Each sequence in the tree is assigned a representative reference sequence label to name the cluster. Any genomes failing the initial quality control (QC) were checked with BUSCO to ensure the genome was a *Campylobacter* species and not contaminated. In addition, simple contig statistics of N50, number of contigs and length of contigs on the assembly were collected. After running through the RC protein workflow (on Terra.bio) the final deduplicated sequences were clustered with MMSeqs2 with parameters for sequence identity at 100% identity and coverage.

Any clusters containing a single genome were checked again at this stage and if the sequence assembly was < 1.6MB in length or contained > 50 contigs the genome was removed. The final number of genomes in this *Campylobacter* cluster reference set was 683.

To look at alternative tree and cluster methods, trees and clustering were carried out with this common set of *Campylobacter* cluster reference 683 database sequences representing ribosomal proteins across the tree of life according to the following protocols:

1. RC protein and MMSeqs2 clustering
2. PopPUNK core genome and PopPUNK clustering
3. The *gyrB* gene sequence and MMSeqs2 clustering

The comparisons of clustering are presented in Figure 3. Initially PopPUNK was used with the poppunk_assign method to their *C. jejuni* provided database and -- update-db. At the time of writing, *C. jejuni* is the only provided *Campylobacter* species database in PopPUNK. The update-db option updates the reference database to map new query sequences (PopPUNK manual). However, we found the use of - - create-db option provided more coverage of our *Campylobacter* cluster reference 683 database sequences.

**Figure 3.**
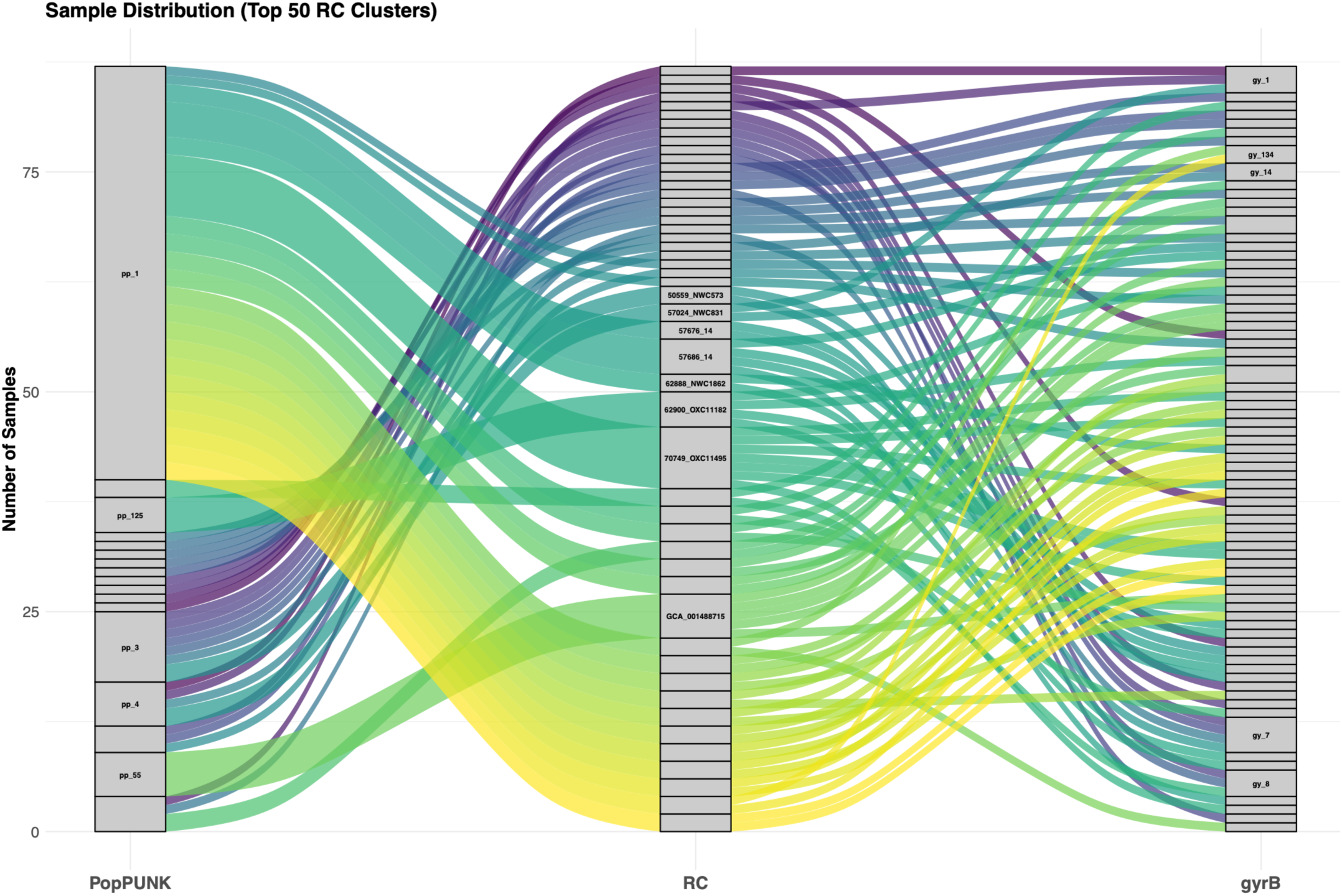
Sankey Alluvial diagram of the top 50 most abundant RC ribosomal protein clusters created from the *Campylobacter* cluster reference 683 database sequences compared to their PopPUNK and *gyrB* clusters

The figure shows the flow of the 683 *Campylobacter* isolate reference genomes tracked through three distinct typing methodologies to illustrate alternative cluster congruence and fragmentation. The three distinct typing methods were used on the identical set of query sequences. For clarity, we have plotted the top 50 most abundant clusters from the RC clustering. Labelled boxes represent the cluster given names for the most abundant clusters from each method. The vertical height of each box corresponds to the total number of samples assigned to that cluster. Connecting ribbons represent the distribution of samples shared between clustering methods. Abbreviations: Genome identifier GCA_001488715 represents GCA_001488715_1_H102740169, 57686_14 represents 57686_14_P0178_03_13, and 57676_14 represents 57676_14_P0124_04_13.

Our results indicate that the RC protein and MMSeqs2 clustering on the *Campylobacter* cluster reference 683 database sequences provide a clustering which is at least equivalent to PopPUNK whilst some RC clusters visibly split the larger PopPUNK clusters in Figure 3 (e.g. see PopPUNK cluster 1, split into several RC clusters. We therefore subsequently used RC typing with MMSeqs2 to cluster 19,663 genomes in the *Campylobacter* Global Dataset [retrieved from public databases and quality controlled October 2022]. Given that the MMSeqs2 parameters used were 100% sequence identity and coverage, each cluster is characterised, for each of the 16 genes, by a single sequence that can be considered as analogous to an allele of an MLST locus, and similarly each cluster can be considered as analogous to an MLST sequence type. Some of these unique sequences were discarded, due to the low quality of the included genomes, according to the stated criteria.

When compared to the BUSCO genes set, we saw slightly different trees when using the defined 39 species (Supplementary Figure S2). Therefore, a phylogenetic tree drawn using ribosomal protein sequences from the *Campylobacter* global dataset was calculated and compared for phylogeny validation with the trees analysing the same genomes built on 1) MLST gene sequences 2) BUSCO core gene sequences, using the BUSCO tree as the gold standard (Supplementary Figure S3).

Using weighted Robinson Foulds (17), Shannon diversity or Faith’s sum of branch lengths as a quantitative way to compare the trees, the RC output (with either DNA or protein sequences) was closest to the gold standard BUSCO tree, giving us confidence in the phylogenies generated by the RC methodology. We included the 7-gene MLST as a well-known typing scheme for reference (Supplementary Figure S3).

To further interrogate the use of the RC proteins we looked at the most divergent members using single ribosomal protein trees – these were RplO, RpsH or RplR. We investigated the variation in these using an alternative measure of conservation: protein structure. On investigation of the AlphaFold database with these protein sequences, we found that the best predicted structure, as determined by local distance difference test (pLDDT), was conserved. It is to be noted that the Eukaryote/Archaeal version of S8 (termed S15A and S22) has an extra secondary structure fold which was less well predicted. We filtered the FoldTree inputs of ribosomal protein S8, S15A and S22 members to include one member of each represented bacterial genus, each member of a set of MAGs and one member of each eukaryotic species. This clearly showed the conservation of the core of this protein across all domains of life. Methods such as FoldTree that create a phylogeny from structural information, a wholly different dataset type, (37) demonstrated a high level of conservation, even in the three most variable proteins, suggesting that the protein (AA) sequence has a conserved core. We therefore believe that clustering based on 16 ribosomal proteins is indeed robust since we expect these proteins to be highly conserved across all life in both sequence and structure. We then generated a Densitree to visualise a consensus tree built using the alternative methods (Supplementary Figure S4). In the Densitree we were able to identify that the MashTree and *gyrB* had a different placing of *C.coli /C.jejuni* and *C. hepaticus* than did all the other trees calculated by different methods. This confirmed RC typing as our method of choice for creation of a reliable phylogeny from the *Camplylobacter* genomes.

Using the cluster reference set of 683 *Campylobacter* genome – generated from the RC typing workflow – we were able to name each of the clusters according to the reference genome for that cluster.

### Stage 3 Investigating isolates from Bangladesh using the RC workflow

For the assignment of unknown genomes to clusters, the RC typing platform generated a phylogeny in Newick format along with intermediate files. Trees were visualised in MicroReact (https://microreact.org) and/or Phandango (https://jameshadfield.github.io/phandango). A phylogenetic tree combining the cluster reference set for *Campylobacter* and Bangladesh strains was constructed. Using RC typing on the Terra platform we clustered 98 novel genomes isolated in Bangladesh and assigned them to 18 different clusters. All clustered with known *C. jejuni* or *C. coli* genomes; seven clusters, however, were novel compared to our reference dataset.

Seven of the 18 clusters identified amongst the 98 Bangladesh isolates lie outside of the *Campylobacter* reference set (683) defined clusters suggesting they are new variants. Further investigation is warranted to determine if these represent different species or just new variants.

## Discussion

Robust Cluster (RC) typing using the Ribosomal Sequences Set on the Terra platform allowed biologists, without specific informatics training, to type *Campylobacter* from Bangladesh to define new clusters.

To analyse the unknown genomes, it was necessary to select a set of 16 *Campylobacter* ribosomal protein sequences (from a reference organism) to create the Ribosomal Sequences Set and then generate a set of clustered reference genomes (Cluster Reference Set) from all publicly available genome sequences (*Campylobacter* Global Dataset). As this required too much compute power to be carried out on Terra, the work was done in the UK using in-house compute power and made available within the compute environment to analyse any set of *Campylobacter* genomes. The method has been made available from Theiagen Genomics via a YouTube video (see code availability section), and the sequences have been made available through multiple websites including jidcuk.org. We hope it may be possible to use a single set of ribosomal protein genes for all bacterial RC typing but this remains to be verified. We also encourage others to generate Cluster Reference Sets for other groups of organisms.

To make sure the typing method was robust, we tested RC against other methods.

The single gene phylogeny using *gyrB* was as good as RC for clustering *Campylobacter* at a sub-species level. The *gyrB* phylogeny was not as close to our gold standard as RC, so we took RC typing forward. This enabled any genome to be related to genomes already deposited and to be designated a species name. Importantly, however, when test isolates did not align with a representative isolate, they were placed in a new cluster and given the name of the test isolate. These will be added to the database of reference cluster genomes as new representative isolates after comprehensive characterisation. The whole process requires careful quality control as contaminated genome sequences or non-*Campylobacter* sequences would falsely increase the apparent diversity. To control for sequencing errors and contamination we included a QC process within the RC program which, as well as normal DNA sequence quality parameters, excluded any genomes with coding sequences of an expected length, the IQR from all available sequences was used (Figure 1). We saw no evidence of ribosomal protein paralogues and so multiple copies in a single genome were excluded as representing contamination.

Our final concern was around introgression of horizontally acquired DNA sequence which is common in the core of *Campylobacter* genomes. Despite the disruption of co-adapted core proteins this appears to be driving niche adaptation in *Campylobacter* (33) and results in hybrid clusters. Whether these would be picked up with the RC method would depend on the involvement of ribosomal proteins and highlights the interest in looking in more detail at new clusters. Although we will have missed some informatic tools we have tried to include the most popular ones used for the typing at species level using whole genome sequence data.

We investigated the use of core SNPs for clustering the *Campylobacter* genomes within the *Campylobacter* global dataset. Whilst there was insufficient diversity in the core genomes across all the *Campylobacter* genus due to low numbers of conserved SNPs, this was also true for ANI. We found calculating AAI to require highly significant compute, and using FastAAI good for quick typing but with insufficient resolution for our purpose (Supplementary Figure S5), the use of core SNPs was useful to sub-divide RC groups, in a way similar to MAC typing for dividing MLST groups of *Salmonella* (21). RC typing supports broader geographic sampling through more accessible analysis of bacterial variation, benefiting global pathogen risk understanding, especially in lower-income countries. This method is being developed for understudied bacteria such as non-aureus staphylococci, *Acinetobacter*, and non-aeruginosa pseudomonads, but it can be used for any organism group.

## Conclusion

Robust Clustering (RC) is a computationally simple, universal typing tool available on Terra.bio. and can be used to analyse genome data from any source, including from portable platforms such as MinION. The RC typing methodology delivered 683 distinct clusters of all *Campylobacter* from publicly available genomes. New clusters were seen when isolates from Bangladesh were tested. Further development to establish Cluster Reference Sets of genomes for other groups of bacteria will extend the utility of this method and will help to address the current bias in genome databases which are overpopulated with genomes of isolates from the developed world. In particular, this method would be useful in bacterial genera where species definition is problematic. We are aware of projects on *Pseudomonas, Staphylococcus* and *Acinetobacter*.

## Online Methods

### Sample collection

All study procedures adhered to the ethical principles of the Declaration of Helsinki and complied with US Institutional Review Board (IRB) guidelines and federal regulations for human subject research (45 CFR 46, the Common Rule).

Ethical approval for the study was obtained from the Ethics Committee of the Child Health Research Foundation (No.BlCH-ERC-0 I /03/2019) and George Mason University (1506712-4). Prior to participation, the study objectives and procedures were explained to all potential participants. Informed consent was obtained from all participants, or from a parent and/or legal guardian, who were informed of their right to withdraw from the study at any time without any consequences.

### Isolation of *Campylobacter*

*Campylobacter* was isolated from the stools of healthy mothers, children and household animals, and was carried out at Kumudini Hospital, Mirzapure, Bangladesh by plating on a selective medium at 37 ^O^C in a microaerophilic environment using anaerobic jars. Identification of suspected *Campylobacter* colonies was by microscopy and confirmed with MinION genomic sequencing.

### Short read DNA Sequencing

DNA extraction with Promega extraction kits was carried out before shipping DNA to the UK for Illumina library prep and DNA sequencing (NextSeq 500).

### *Campylobacter* genomes from publicly available sequence databases

For 39 representative genomes described in (32), BUSCO v5 (38) was executed and completeness of each genome assessed. Not all BUSCO genes were detected in all genomes. The full downloaded dataset was first analysed for outliers of assembly statistics (total genome length, number of contigs, N50, average contig length, length of longest contig and length of the shortest contig) and secondly by multi-dimensional scaling of Mash (39) pairwise distance matrices, following the workflow proposed for QC in the Panaroo clustering tool (40). Outliers were subjected to examination using BUSCO v5 to check for the presence of the *Campylobacter* lineage set of housekeeping genes. Genomes were removed if considered to be less than 100% complete.

### Genomic methods for assigning species

To test the ability to differentiate *Campylobacter* species, each typing scheme was assessed on 39 published genomes and only taken forward if 39 species were identified. Testing was carried out using software installed on the MacBook Pro M2 (2023) with arm64 architecture running OSX 13.6.3 with 64Gb memory using 12 threads (Results in Table). Where dependency libraries used by the software under test were incompatible with arm architecture, OSX was used with Rosetta 2 ahead of time compilation to emulate x86_64. For processes running in under 5 minutes, results are the mean of 10 runs with a warm-up run; longer running processes were timed once. We do not rule out the possibility that runs were memory limited in some cases which may have led to longer runtimes.

### Single Nucleotide Polymorphisms (core SNPs)

A core SNP method was included because it is used by centralised public health laboratories (e.g. UKHSA SnapperDB (41)) for outbreak investigations. Here we used the snippy v4.0 pipeline (42).

### 16S gene phylogeny

The naming of bacterial species using 16S sequence is recognised by international committees and was included here (Supplementary Figure S1). *Campylobacter* genomes usually have three copies of the 16S gene so all three copies were extracted from each of the 39 draft genome assemblies. These were aligned with Mafft v7.49 and a phylogenetic tree created with Iqtree v2 (43) (Supplementary Figure S1).

### Single gene gyrase B subunit, *gyrB*

The *gyrB* DNA sequences were identified from each assembly using BLAST v2.4.1 and extracted from the BLAST XML file. These were aligned with Mafft and phylogenies created with Iqtree.

### Hug nucleotide & Hug protein (RC)

The phylogenies were constructed using the snakemake 8.10.8 workflow management system as our RC snakemake (35) pipeline with Python (Figure 1). Parameters are available to choose either the nucleotide or the protein sequences, depending on requirements and the level of conservation in the genomes to be tested. Phylogenies presented in this paper were built using the web-based Terra.bio system. Quality control was built into the full workflow, for example, carried out on the ribosomal protein sequences of the *Campylobacter* global sequence set, using variation in the length of the ribosomal proteins annotated as such (Figure 1) and so for inclusion of ∼95% of proteins the predicted CDS had to be within 1.5 x the IQR from the median for all sequences from all organisms. We generated the Ribosomal Sequences Set from a well-known *Campylobacter* genome (NCTC 11168). In this way we hope that the pipeline can be used for genomes from any group of *Campylobacter spp*.

### BUSCO phylogenies

All the Pfam v36.0 hidden Markov models for the genes identified as core by BUSCO v5.5 throughout the bacterial kingdom were gathered into a single HMM file. This was used in HMMER v3.4 hmmsearch to provide a table containing hits for each sample tested. Gene names identified as hits were used to extract the protein sequences from the GFF3 files. The extracted protein sequences were concatenated together on a per strain basis. These concatenated sequences were used in Mafft alignments and IQtree was used to create phylogenies.

### Mash

The Mash (v2.2.2) approach is based on kmers and is excellent for speed and data-intensive problems and generates an estimation of the mutation rate to rapidly generate approximate phylogenies using hierarchical clustering. Mash is based on MinHash sketches and incorporates a comparison of these sketches to provide an estimate of the Jaccard index (fraction of shared kmers). Due to limitations of the kmer approach and the simple distance model, the original authors emphasise that Mash is not explicitly designed for phylogenetic reconstructions, especially those with high divergence or large size differences.

A more recent addition, Mashtree (v1.4.6) (44), was used in this study which creates dendrograms from Mash distances and was executed with 100 bootstrap replicates on 12 threads directly from DNA sequence FASTA data at a minimum depth parameter of 0.

### CheckM

CheckM v1.1.6 is a lineage identifier and phylogenetic placer written in Skala with the aim of investigating metagenomic bins. It is currently based on hidden Markov model searches against a database of marker genes. CheckM is a powerful and useful phylogenetic placer of metagenomic bins. CheckM requires a specific archived version of HMMER (v3.1), a protected container environment running older software. The marker database is from 2015 and has nonetheless relatively large memory and compute requirements. Whilst CheckM placed the 39 groups separately on a marker tree, however, for the lineage estimation, the workflow provided two lineages within the *Campylobacter* genus and the other 37 were assigned to Epsilonproteobacteria.

### Panaroo

Panaroo v1.5.0 was used to calculate the pangenome from a provided set of genomes. The process does not build a phylogenetic tree but rather a network.

### PopPUNK

PopPUNK (45) is a method designed to analyse new query sequences and clusters of genomes from existing genome databases. PopPUNK v2.7.0 was executed according to the documentation using PopPUNK assign and using the available *Campylobacter jejuni* database. We also tested the clustering tool in PopPUNK to analyse the output from the RC pipeline.

### Circos and Tanglegrams

Scatterplots and tanglegrams were carried out in R programming language v4.3.2. The Sankey alluvial diagram was carried out using ggalluvial and ggplot2 in R v4.4.3.

### Generating a cluster reference set

We were able to generate a representative set of *Campylobacter* genomes as a cluster reference set. After clustering was carried out using MMSeqs2, with 100% identity, quality control was performed by first identifying the quality of each genome assembly in terms of total sequence length, number of contigs and N50. These statistics were then compared on each of the sequences found per cluster of the ribosomal proteins. For each cluster the best representative sequence was then chosen, based on the assembly statistics. If more than one assembly was excellent, the longest assembly was chosen. If only a single representative was found in a cluster, and the assembly was poor based on the statistics when examined using the Panaroo QC pipeline, the cluster was deleted.

### Visualisation, clustering programs and statistical comparisons DensiTree

DensiTree v3.0.3 is a program written in Java for qualitative analysis of sets of trees (46–48). Using Densitree we were able to visualise a consensus tree built using several alternative methods (Supplementary Figure S4).

### FoldTree

Foldtree v0.0.1 was executed as described in the results section and produced a tree of RpsH protein folds across all kingdoms, as shown in Supplementary Figure S3.

### Bayesian hierarchical clustering

We evaluated but did not use RHierBAPS (49) and FastBAPS (50) algorithms to apply hierarchical Bayesian clustering to nucleotide genetic sequences using the same likelihood as BAPS (51). Whilst BAPS clustering with FastBAPS is designed to be generated ‘on the fly’, here we seek to create a stable clustering for the population structure of *Campylobacter* using a dataset of *Campylobacter* genomes from both pubMLST and GenBank (*Campylobacter* global dataset). We evaluated BAPS clustering for *Campylobacter* within our RC workflow with either protein or DNA sequences. However, protein sequence clustering is not supported by FastBAPS. We produced functions written in R for use with protein sequences for RhierBAPS [see code availability section], however, since BAPS clustering depends to an extent on the number of sequences in a cluster and it is not possible to create a cluster with one representative, a solution with more stable clustering was sought.

### MMSeqs2 clustering

MMSeqs2, an ultrafast clustering program from (36) was used with the easy-cluster option and parameters for exact identity match (-c 1.0).

### Generation of a reference set

Public databases pubMLST (as BIGSdb) and GenBank/EMBL were searched for *Campylobacter* genomes to create a dataset of 19,663 genomes [Accessed October 2022]. We checked the quality of all 19663 genome sequence assemblies using the Panaroo quality control guidelines. Assemblies were processed with Mash distances, then multi-dimensional scaling (MDS).

Alignment and tree building methods

Alignments were built with Mafft v7.490 and trees with Iqtree2 v2.3.6

Robinson Fould index and the Shannon index

To compare the trees generated against our gold standard, BUSCO tree, we used the Robinson Fould and Shannon indexes (R packageTreeDist 2.12.0).

### Clarification of dataset nomenclature

Ribosomal sequences set (RCprot) = the 16 proteins used for each genus selected from a designated reference strain

Genus specific Cluster Reference Set = the 683 unique syntenic sequences taken from the *Campylobacter* Global Dataset of *Campylobacter* genome sequences and used to represent and define all the currently described ribosomal clusters within the *Campylobacter* genus

*Campylobacter* Global Dataset = all publicly available 19,663 genome sequences from organisms annotated as *Campylobacter*, as at October 2022.

RC is used to denote the typing method – Robust Clustering. RC sequences may be nucleotide or protein.

## Acknowledgements

We acknowledge the contributions of Jibon Hossain, Rasel Mahmud, and Nusrat Alam during culture, isolation, identification and sequencing of *Campylobacter* organisms in Bangladesh. The QIB Bioinformatics team supported access to data from in-house sequencing and The QIB sequencing team sequenced the DNA from Bangladesh isolates. Theiagen Genomics implemented the software package on the Terra platform, and we continue to further develop this package with them on a contract research basis. Caroline Jarvis proofread the final document and Anne Priestly (Freelance academic editor) corrected the English in the final version. We also acknowledge Rebecca Clifford (now a GP in York) for her work on the RC typing of *Staphylococcus* that paved the way for this current research, Aidan Wain (freelance) for IT support and Leonardo de Oliveira Martins (Liverpool)

## Funding

LC, JIDCuk Charitable Trust

JW, CH, HF, NJ, BD, were supported by the Biotechnology and Biological Sciences Research Council (BBSRC) Institute Strategic Programme Microbes in the Food Chain BB/R012504/1

AR, George Washington University,

SS, AMT, ns and HR were funded by CHRF, Dhaka, Bangladesh

The collection and sequencing of *Campylobacter* strains in Bangladesh was partially funded by the Bill and Melinda Gates Foundation.

## Author Contributions

JW devised the method and wrote the first draft of the manuscript; LC, developed the script for the original workflow, generated the first reference set of genome sequences from all publicly available genomes, developed the QC algorithms for the Terra platform and edited the manuscript throughout; AMT, NS, HR were responsible for the day to day management and running of the project in Bangladesh, verified the identity of the isolates, extracted DNA in Bangladesh, and commented on the manuscript; CH carried out computational trouble shooting, liaised with Theiagen, refined the Terra workflow; HF established the original workflow for robust cluster typing; NJ managed the *Campylobacter* project from which the isolates came, oversaw the sequencing of isolates and edited the final manuscript; BD was instrumental in setting up the project in Bangladesh and analysed the sequence data; AR was the PI on the original Bangladesh study and edited the final manuscript; SS was responsible for the original design and management of the project, selected the laboratory methods for the isolation of *Campylobacter* and commented on the manuscript.

## Data Availability statement

The final workflow and instructions for use are included in the data.

The Bangladesh *Campylobacter* genome data for this study have been deposited in the European Nucleotide Archive (ENA) at EMBL-EBI under accession number PRJEB96283. The RC workflow can be accessed at Terra.bio using JIDC Dockstore https://dockstore.org/workflows/github.com/jidc-trust/ribosomal-wdl/RCTyping:master?tab=info following the documentation manual presented in the Supplementary Information. The reference dataset can be accessed from Zenodo doi://10.5281/zenodo.15189883 (or https://github.com/LCrossman).

## Code availability

https://dockstore.org/workflows/github.com/jidc-trust/ribosomal-wdl/RCTyping:master?tab=info and https://github.com/LCrossman

