## Supplementary_data for "Robust Cluster Typing, a computationally accessible tool for species designation from whole genome data, exemplified with the genus *Campylobacter*"

Supplementary Figures

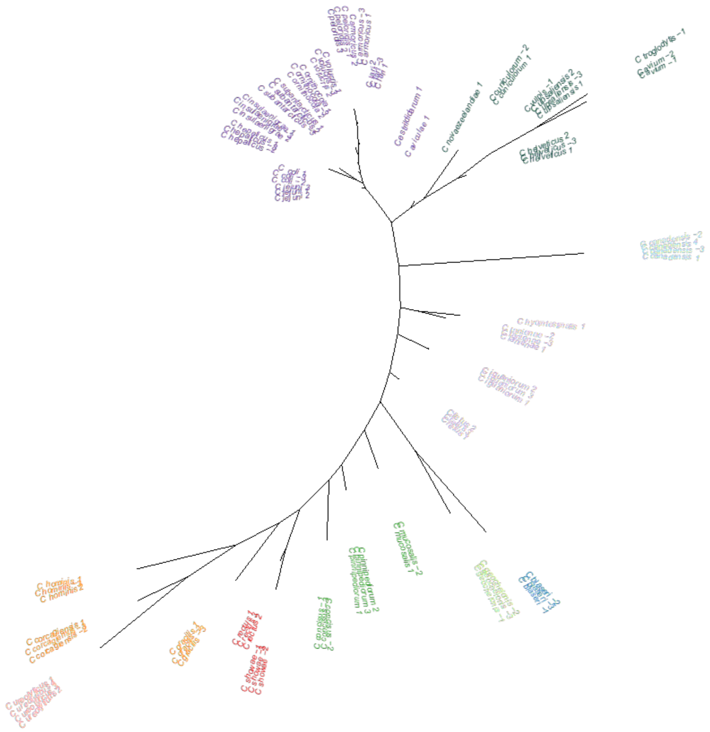

**Suplementary Figure 1** 16S phylogeny including all copies present in the genome

The 16S rRNA genes are named by species and according to the copy number encountered in the genome 1-3. The 16S tree is coloured according to the cluster assigned with FastBAPS v1.0.8 and presented as an unrooted tree. (Tonkin-Hill,G., Lees,J.A., Bentley,S.D., Frost,S.D.W. and Corander,J. (2019) Fast hierarchical Bayesian analysis of population structure. Nucleic Acids Res., 10.1093/nar/gkz361.)

(i) Tanglegram RC protein_v_BUSCO

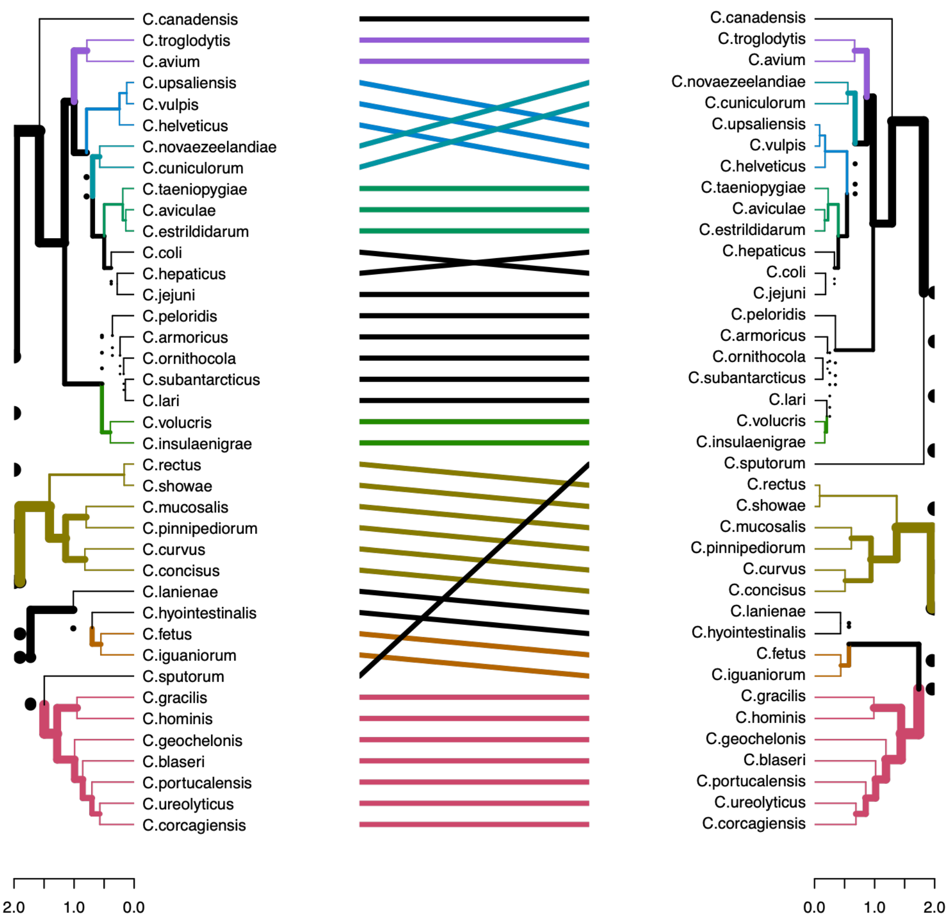

(ii) Tanglegram of Busco vs *gyrB*

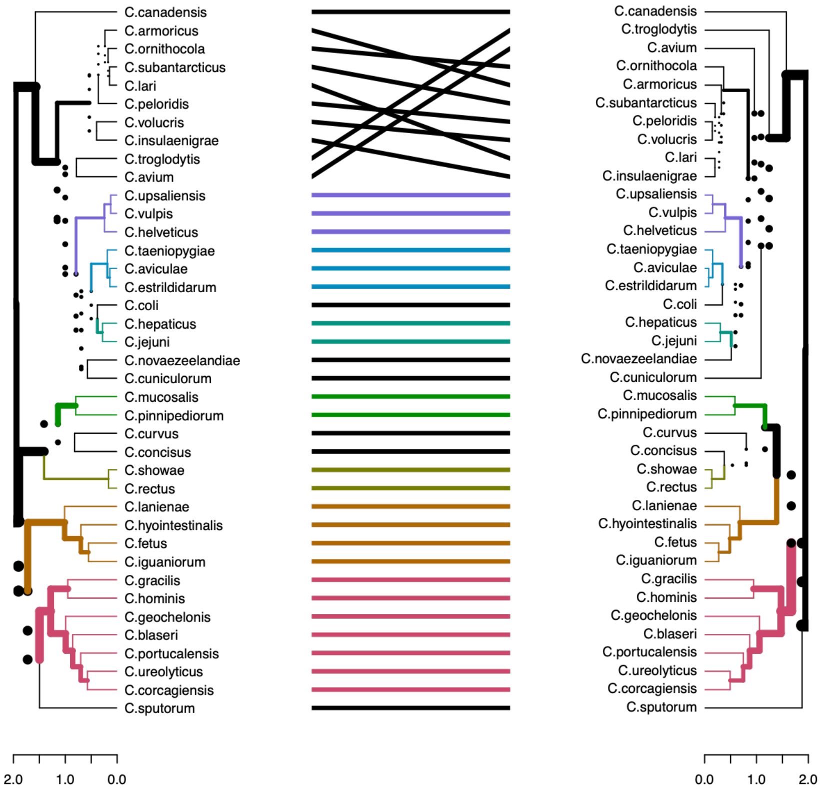


**Supplementary Figure S2**

Tanglegrams comparing alternative trees

Damien M de Vienne, Tanglegrams Are Misleading for Visual Evaluation of Tree Congruence, Molecular Biology and Evolution, Volume 36, Issue 1, January 2019, Pages 174–176, https://doi.org/10.1093/molbev/msy196

Here we provide a set of tanglegrams to aid the comparison of tree visualization. Whilst tanglegrams have been described by de Vienne *et al.* (2019) as misleading for visual evaluation of tree congruence, we have applied the author's suggested best practices which are to include the use of colour and line style to highlight the similarities and differences between the compared trees. (i) RC protein tree on the left hand side compared to BUSCO tree on the right hand side (ii) BUSCO tree on the left hand side compared to gyrB tree on the right hand side.

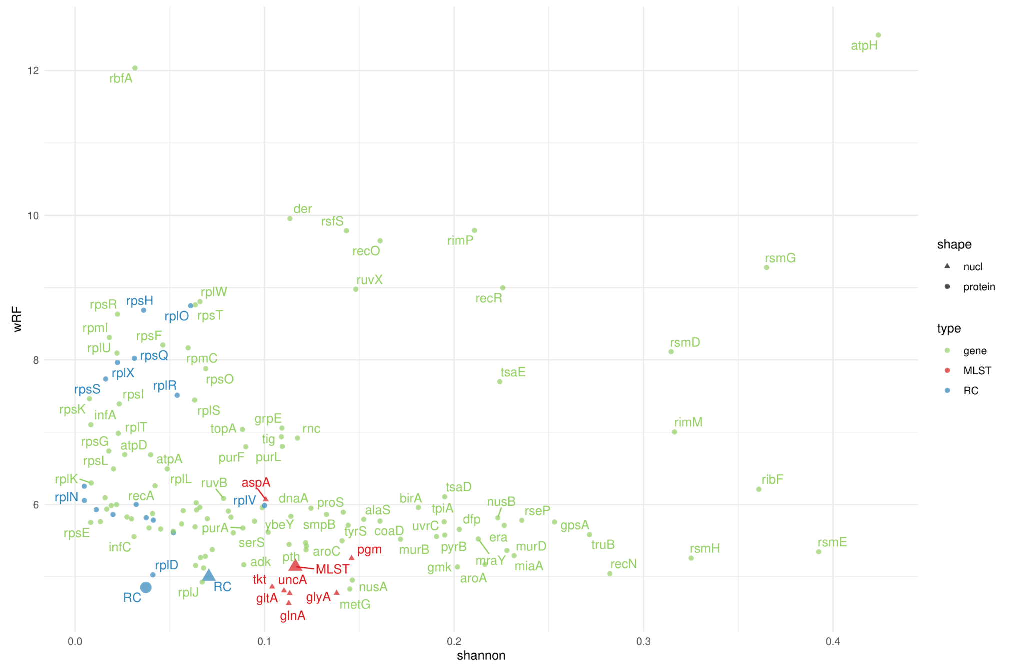


FigS3 A

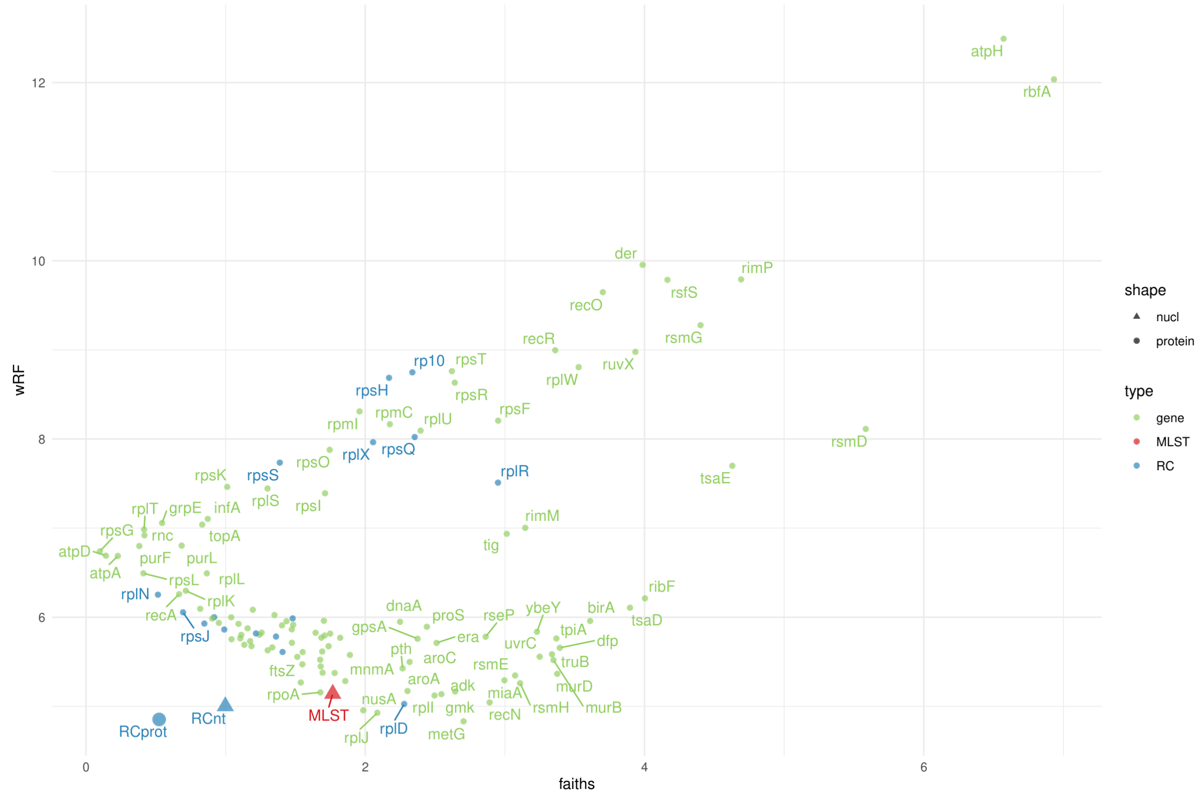


FigS3 B

**Supplementary Figure S3**.

Scatterplots in R showing the within group diversity against the weighted Robinson Foulds comparing with a gold standard tree of concatenated BUSCO proteins. Shown in green and marked with the name are the statistics from alignments and trees of orthologues, from at least 19,658 genomes, of that separate gene (all gene sequnces passing QC were included). For example, marked in Fig 3A, *ybeY* is plotted as the Shannon diversity of the alignment of 19,659 orthologues of *ybeY* from the genome dataset on the x axis and the weighted Robinson Foulds statistic comparing the *ybeY* tree against the whole BUSCO tree.

We also considered the MLST genes for *Campylobacter jejuni* and *coli*, which are *aspA, tkt, gltA, glyA, pgm, glnA* and *uncA*. Each of these statistics are plotted in red, alongside the statistics for the whole MLST tree. Genes marked in blue are those that are part of the RC typing scheme, we also include the Shannon diversity and weighted Robinson Foulds for the concatenated RC protein alignments and tree and RC nucleotide alignments and tree. For clarity statistics using nucleotide data are shown as a triangle, and those with protein data are shown as a circle.

**Supplementary Figure 3A**:
We calculated the Shannon diversity on each of the separate orthologue alignments to plot on the x axis, this indicates the diversity within each alignment, plotting the weighted Robinson Foulds of each separate tree in comparison to a gold standard of a phylogenetic tree from concatenated BUSCO proteins for *Campylobacter* across the full dataset.

**Supplementary Figure 3B:**
We calculated the Faiths sum of branch lengths, the phylogenetic diversity, from each of the separate orthologue trees and plotted against the weighted Robinson Foulds of each separate tree in comparison to a gold standard of a phylogenetic tree from all the concatenated BUSCO proteins for *Campylobacter* across the full dataset.

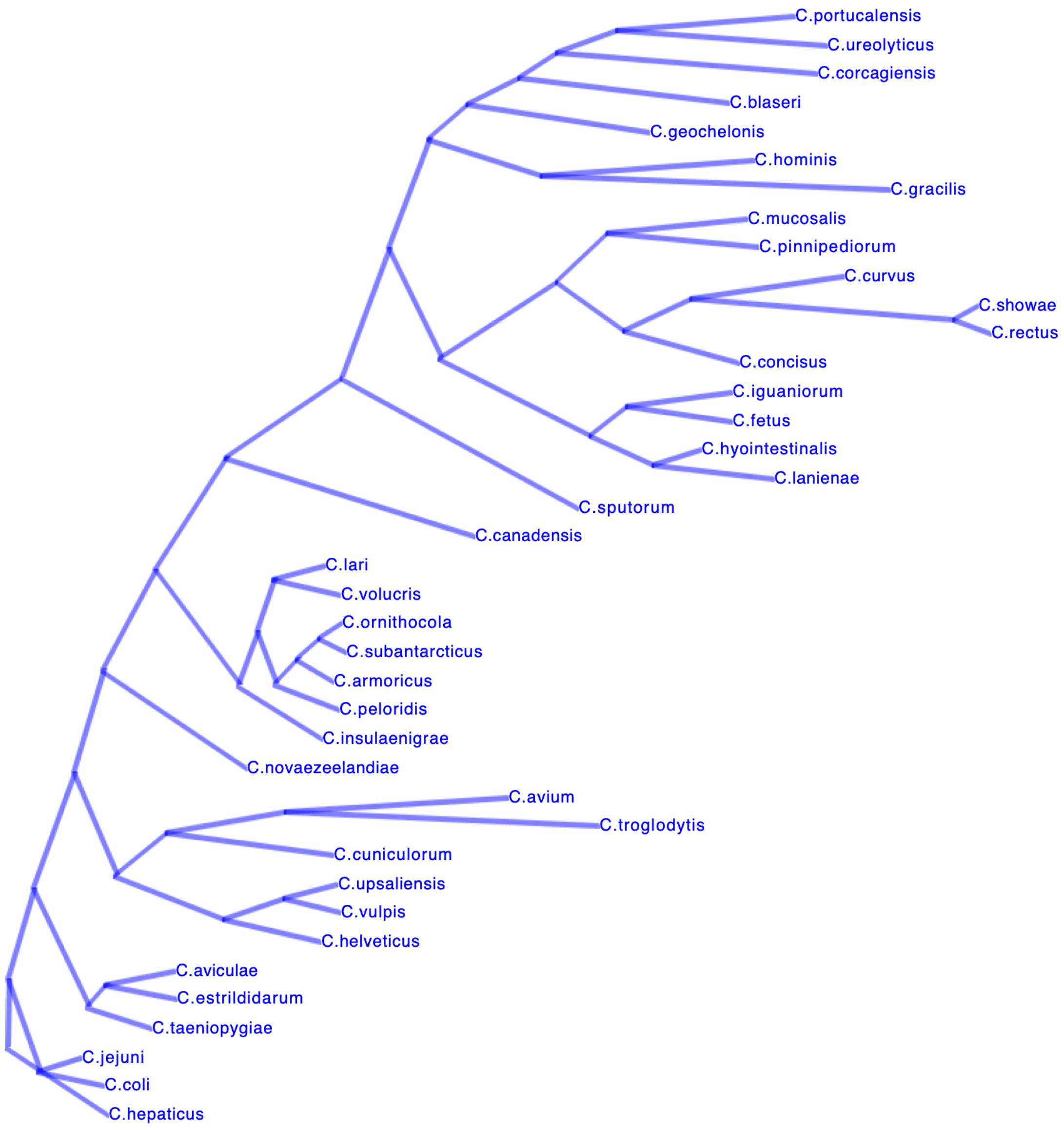


**Supplementary Figure S4.** DensiTree Visualization – A joint tree composed from different methods of the 39.

This Densitree is a consensus phylogeny composed of trees built by the alternative methods described in the methods for the 39 species dataset. Comparing this figure with the *gyrB* and MashTree figures, indicated that they did not fully agree with the other methods in the placing of jejuni/coli & hepaticus. Remco R. Bouckaert, DensiTree: making sense of sets of phylogenetic trees, Bioinformatics, Volume 26, Issue 10, May 2010, Pages 1372–1373, https://doi.org/10.1093/bioinformatics/btq110

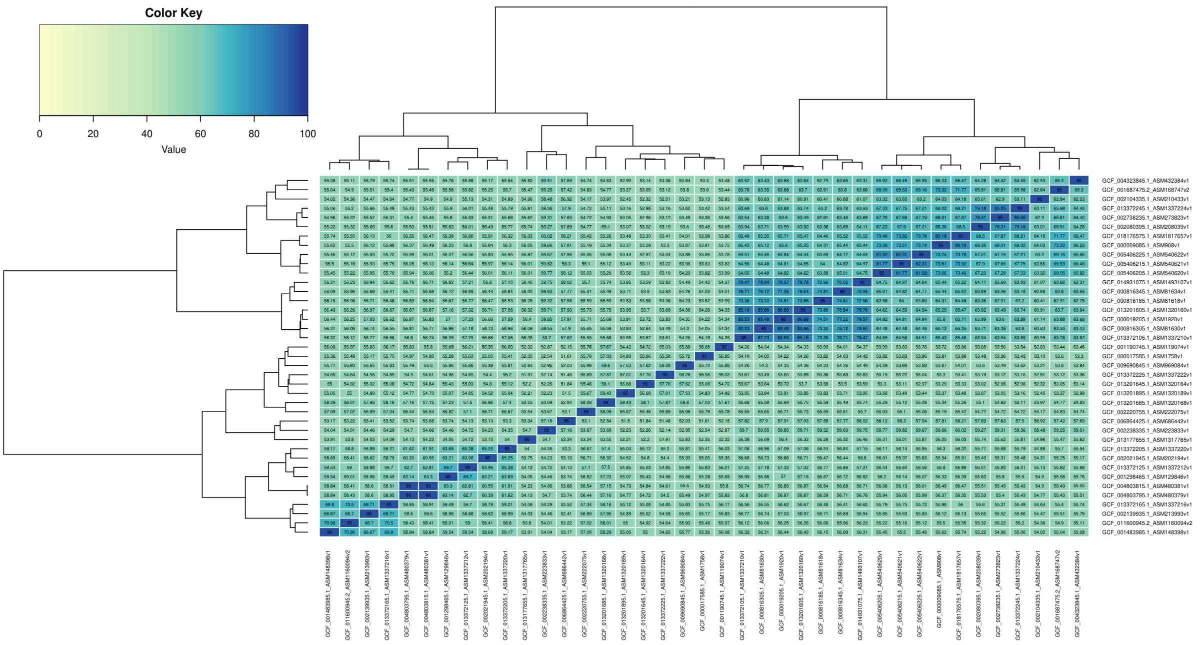


**Supplementary Figure S5**: A heatmap showing the similarirty between the concatonated ribosomal dataset for the 39 desecibed spaicies of Campylobacter using fastAAI. The algorythm is useful for quick comaprisons of disperate bacteria but, unfortunately gives less than 100% concordance when the same genome is compared. It was therefore not used in this study.
